# Interpreting ruminant specific conserved non-coding elements by developmental gene regulatory network

**DOI:** 10.1101/2021.11.08.467829

**Authors:** Xiangyu Pan, Zhaoxia Ma, Xinqi Sun, Hui Li, Tingting Zhang, Chen Zhao, Nini Wang, Rasmus Heller, Wing Hung Wong, Wen Wang, Yu Jiang, Yong Wang

**Author notes:** These authors contributed equally to this work (Xiangyu Pan, Zhaoxia Ma, Xinqi Sun, Hui Li). Corresponding authors. (Y.W.), (Y.J.), (W.W.).

## Abstract

**Background:** Biologists long recognized that the genetic information encoded in DNA leads to trait innovation via gene regulatory network (GRN) in development.

**Results:** Here, we generated paired expression and chromatin accessibility data during rumen and esophagus development in sheep and revealed 1,601 active ruminant-specific conserved non-coding elements (active-RSCNEs). To interpret the function of these active-RSCNEs, we developed a Conserved Non-coding Element interpretation method by gene Regulatory network (CNEReg) to define toolkit transcription factors (TTF) and model its regulation on rumen specific gene via batteries of active-RSCNEs during development. Our developmental GRN reveals 18 TTFs and 313 active-RSCNEs regulating the functional modules of the rumen and identifies OTX1, SOX21, HOXC8, SOX2, TP63, PPARG and 16 active-RSCNEs that functionally distinguish the rumen from the esophagus.

**Conclusions:** We argue that CNEReg is an attractive systematic approach to integrate evo-devo concepts with omics data to understand how gene regulation evolves and shapes complex traits.

## Background

To answer the key question of how new traits arise during the macro-evolutionary process, biologists have long realized the necessity to understand the gene regulation in development responsible for morphological diversity, i.e., which genes are expressed, what regulatory element changes are involved and how do regulatory element changes affect development [1]. Only recently has the field of large-scale omics and the accumulation of data matured sufficiently to explore these theoretical concepts in detail. Here, we investigate the ruminant multi-chambered stomach, a key mammalian organ innovation and a cornerstone of evolutionary theory, as an example to illustrate a novel framework for integrating multi-omics data to address the fundamental question of organ innovation.

The rumen hosts a diverse ecosystem of microorganisms and facilitates efficient plant fibers digestion and short chain fatty acids uptake, which significantly promoted the expansion and diversification of ruminant animals by providing a unique evolutionary advantage relative to non-ruminants [2]. This remarkable morphological innovation raises the fundamental question of how the genetic toolkit generates functional complexity through development and evolution [1, 3, 4]. By comparing 51 ruminants with 12 mammalian outgroup species genomes, we previously identified 221,166 ruminant-specific conserved non-coding elements (RSCNEs), which span about 0.61% of the genome (16.5 Mbp in total) [5]. These RSCNEs are potential regulatory elements of proximal or distal genes for transcriptional regulation in the development of morphological and physiological traits [6]. In addition, we previously sequenced two representative ruminants (sheep and roe deer) for gene expression across 50 tissues. Comparative transcriptome analysis reveals 656 rumen-specific expressed genes (RSEGs) and hypothesizes that rumen’s anatomical predecessor is the esophagus by their most similar expression profile [5, 7]. It’s in pressing need to understand how the RSCNEs leading to the expression of RSEG changes.

One major bottleneck is that the cellular context, target gene and mode of gene regulation of the RSCNEs are largely unknown. First, the regulatory role of RSCNEs could be spatio-temporally dynamic and highly context-specific. Second, some RSCNEs were located distant (e.g., more than 500 kbp) from any genes and therefore could not be associated with any target genes using standard approaches, such as GREAT [8]. This problem is emphasized by a recent finding that non-coding region associating with a human craniofacial disorder causally affects SOX9 expression at a distance up to 1.45 Mbp during a restricted time window of facial progenitor development [9]. This example motivated us to develop a framework for RSCNE functional inference by uncovering GRNs at different developmental times and in different tissue types, and integrating them with their functional relation to traits.

To tackle the above challenges, we generated time series of paired gene expression and chromatin accessibility data during rumen and esophagus development in sheep to reconstruct a time series of developmental GRNs. Our previous efforts showed that jointly modeling multi-omics data allows us to infer high quality tissue specific regulatory networks [10], which can be used to identify key transcription factors (TFs) during differentiation [11], reveal causal regulations [12], and interpret functionally important genetic variants [13]. Taken together, we aim to integrate multi-omics data to a reconstruct genome-wide GRN during different stages of development in an apomorphic organ. Specifically, this allows us to understand how transcription factors bind to functional RSCNEs to coordinate cell type specific gene expression of rumen-specifically expressed genes (RSEGs), and hence to gain further insights into the evolutionary development of new organs.

## Results

### Landscape of accessible chromatin regions and gene expression during rumen development

We resolved a high-resolution chromatin accessibility and gene expression landscapes of rumen development by collecting ruminal epithelial cell, esophageal epithelial cell, and hepatocyte cell at five stages (embryo 60-day [E60], postnatal day 1 [D1], day 7 [D7], day 28 [D28] and adult 1-year [Y1]) from 14 sheep (Fig. 1A). Our experimental design covers the major stages of the ruminal epithelium differentiation and development [14, 15], and ensures an exact matching of tissues used for RNA-seq and ATAC-seq libraries. In total, 37 ATAC-seq and 34 RNA-seq data sets including biological and technical replicates showed high quality (Methods; Additional file 1: Table S1, 2). The ATAC-seq samples have an average of 115 Mbp post-quality control uniquely mapped fragments to the sheep Oar_4.0 genome (Additional file 1: Table S1; Additional file 2: Fig. S1A), which are highly enriched at transcription start sites (Additional file 2: Fig. S1B) and show a nucleosome structure consistent distribution (Additional file 2: Fig. S1C). We obtained 178,651 open chromatin regions (OCRs) across all samples (mean 46,872 peaks per sample) (Additional file 1: Table S1).

**Fig. 1.**
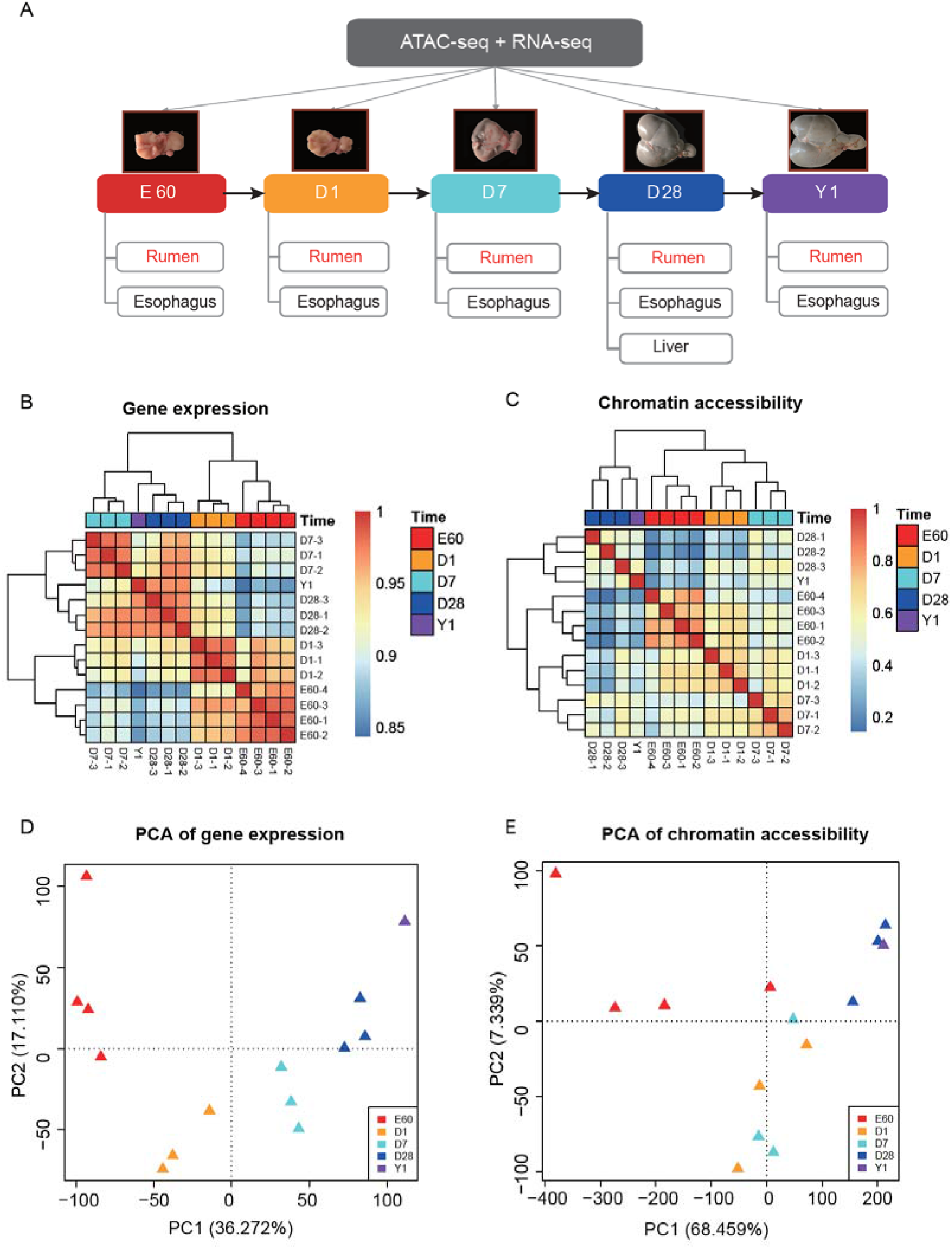
Paired expression and chromatin accessibility time series data reveals regulatory landscape for rumen development. **(A)** Experimental design diagram for multi-replicate, multi-tissue, and multi-level omics data profiling during sheep development from embryo 60-day (E60), postnatal day 1, 7, 28(D1, D7, and D28) to adult 1-year (Y1). **(B, C)** Hierarchical clustering of gene expression for 14,637 genes and chromatin accessibility for 178,651 open chromatin regions both indicate rumen’s multi-stage development process. D28 and Y1 are more closely grouped and E60, D1, and D7 are distinct group both in expression and chromatin accessibility. **(D, E)** Unsupervised principal component analysis of rumen’s gene expression and chromatin accessibility. Multi-stage development pattern is consistent with clustering results. Early development stages E60 and D1 show larger replicate variation than D7, D28, and Y1 at both chromatin accessibility and gene expression. In addition, chromatin accessibility shows more smoothed trajectory than expression.

Hierarchical clustering of gene expression and chromatin accessibility show that rumen development is a multi-stage biological process (Fig. 1B, 1C). Stages E60 and D1 cluster in one group and D7, D28, and Y1 cluster in another group by gene expression. Chromatin accessibility patterns further distinguish stages E60 and D1. Principal component analysis (PCA) for 14,637 expressed genes and 178,651 OCRs corroborates this multi-stage pattern (Fig. 1D, 1E). Early development stages E60 and D1 show larger replicate variation than D7, D28, and Y1 at both chromatin accessibility and gene expression levels (Fig. 1D, 1E). In addition, chromatin accessibility shows a more smoothed trajectory than gene expression during rumen development (Fig. 1C).

The esophagus shows a very similar multi-stage development (Additional file 2: Fig. S2A, B). PCA indicates larger variance in developmental stages (PC1 32%) and smaller variance among tissue types (PC2 25%) (Additional file 2: Fig. S2C, D). This pattern is consistent with previous studies showing that gene expression divergence between tissues/cell types increases as development progresses [16]. Importantly, our chromatin accessibility data mirror this pattern, i.e., the similarity in chromatin accessibility distribution between the two tissues declines as development progresses.

### Identification and characterization of active-RSCNEs

We obtained 159,837 reproducible OCRs by intersecting peaks from three replicates for rumen and esophagus at four developmental stages. The number of reproducible OCRs was largest in stage E60 (about 40%) and decreased along the developmental stages (Fig. 2A), which is consistent with the observation of higher amounts of accessible chromatin in embryonic stage [17]. Most reproducible OCRs were located at distal intergenic (39.42%), intron (32.61%), and promoter (21.46%) (+/−3 kbp from transcription start site) sites (Fig. 2B). After overlapping the OCRs with 221,166 RSCNEs from ruminant comparative genomics analysis [5], we identified 1,601 active-RSCNEs with an average length of 82 bp (Additional file 1: Table S3). Again, the number of active RSCNEs decreases along the development stages both in rumen and esophagus (Fig. 2C). They are mainly located in distal intergenic (48.95%), intron (42.4%), and promoter regions (4.96%) (Fig. 2D). Compared to all reproducible OCRs, active-RSCNEs are less in promoter regions by 15% (Additional file 2: Fig. S3A), and the esophagus shows a consistent trend (Additional file 2: Fig. S3B). This suggests active-RSCNEs tend to function as distal element during development. In addition, our observation that the vast majority of active-RSCNEs are found in early developmental stages (>90% in E60, D1, D7) emphasizes the importance of early developmental cellular context for interpreting the regulatory role of CNEs.

**Fig. 2.**
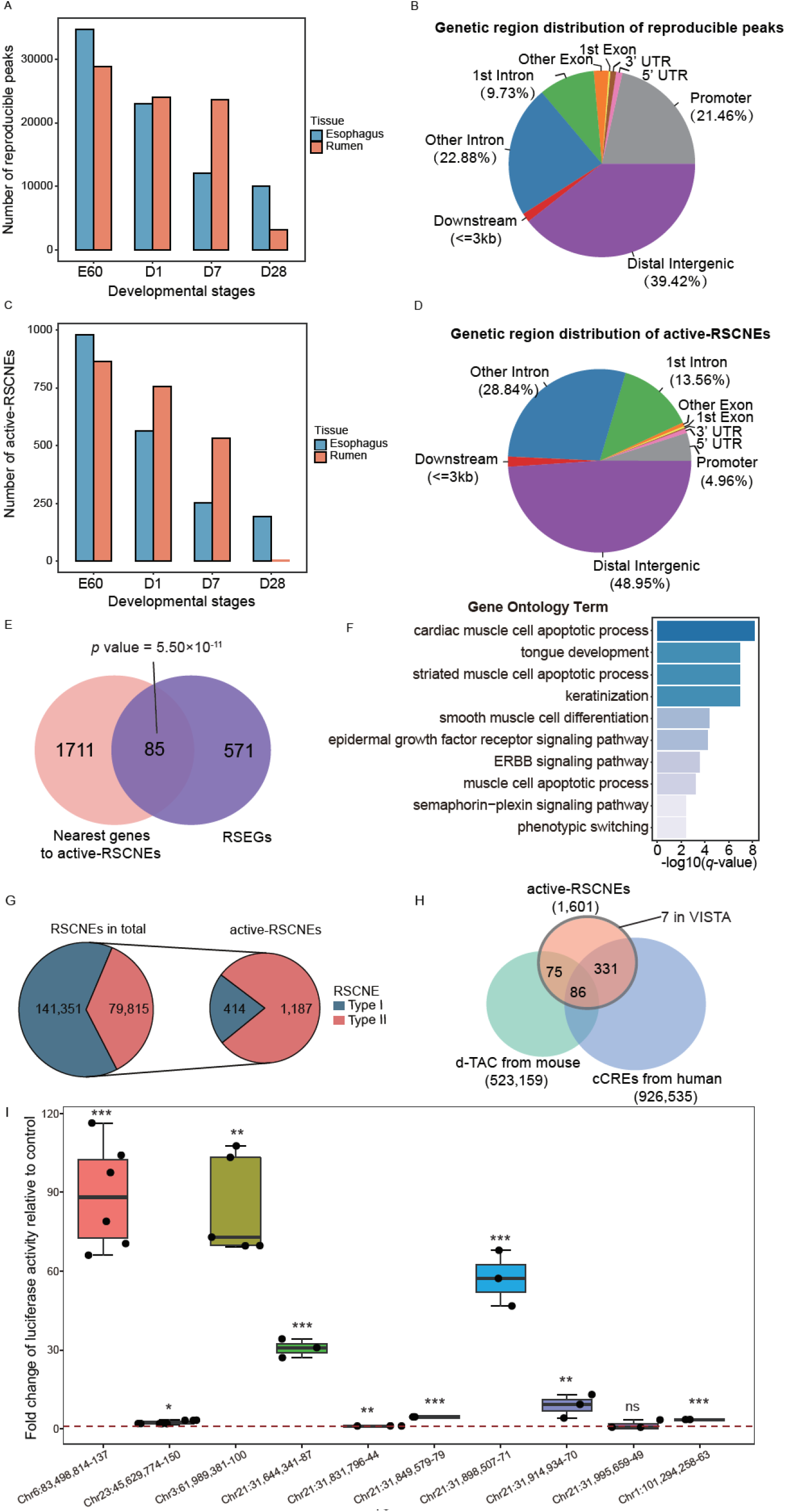
Characterization of active-RSCNEs. (**A**) The number of reproducible peaks during each developmental stage in rumen and esophagus. (**B**) Annotating reproducible peaks by location in different genomic regions. (**C**) The number of active-RSCNEs during each developmental stage in rumen and esophagus. (**D**) Annotating active-RSCNEs by location in different genomic regions. (**E**) GO enrichment analysis for genes near the active -RSCNEs. (**F**) The genes nearest to active-RSCNEs are enriched in RSEGs. *p*-value is calculated by Fisher’s exact test. (**G**) Illustration of the number of Type I and Type II in total RSCNEs and active-RSCNEs. (**H**) The intersections among active-RSCNEs with enhancers from d-TAC, cCREs, and VISTA datasets. (**I**) Luciferase activity assays of 10 active-RSCNEs randomly chosen in 1,601 active-RSNCEs. 9 of 10 show regulatory activity in PGL-3 promoter.

We next associated the 1,601 active-RSCNEs with their 1,796 genes nearby. Gene ontology analysis of these genes are enriched in terms, such as “primary metabolic process”, “catalytic activity”, and “regulation of signaling” (Additional file 2: Fig. S3C). Moreover, those 1,796 genes are significantly enriched in transcription factors (TFs) (Additional file 2: Fig. S3D; Fisher’s exact test, *P* value = 4.20×10^−4^). These 1,796 genes overlap with 656 RSEGs by 85 genes (Fig. 2E; Fisher’s exact test, *P* value = 5.50×10^−11^) which are enriched in “cardiac muscle cell apoptotic process”, “tongue development”, and “keratinization” (Fig. 2F).

The 1,601 active-RSCNEs are composed of 414 Type I and 1,187 Type II RSCNEs (Additional file 1: Table S3; Fig. 2G). Type I have no known orthologs in non-ruminant outgroups and Type II orthologs exhibit significantly higher substitution rates among outgroups [5]. The ratio between Type I and Type II active-RSCNEs is ~0.35, which is 5-fold less than that of all RSCNEs, which have a Type I/Type II ratio ~1.77 (Fig. 2G). This surprising fact suggests that Type II RSCNEs tend to be more activate in the developmental stage than Type I. Because of the deeper evolutionary origin of Type II RSCNEs, they are more likely to function by altering existing regulatory elements. Furthermore, we found that active-RSCNEs are enriched for binding motifs of transcriptional regulators known to play a vital role in rumen development (AP-1, PITX1, TP63, KLF, GRHL, TEAD, OTX, and HOX) (128 motifs with Benjamini *q*-value <1.00×10^−3^ are listed in Additional file 1: Table S4), suggesting that some active-RSCNEs may act as rumen developmental enhancers.

To assess whether the RSCNEs are likely to play an enhancer role, we next compared our 1,601 active-RSCNEs with the 523,159 developmental regions of transposase-accessible chromatin (d-TACs) data sets from mouse [18] and 926,535 human enhancers from ENCODE phase III [19]. About 24% of the active-RSCNEs can be found in these data sets (Fig. 2H), and 11 active-RSCNEs show *in vivo* reporter activity according to the VISTA database [20] (Fig. 2H). To validate the potential regulatory activity, 10 active-RSCNEs of length ~300 bp were randomly selected and assessed for enhancer activity detection in both sheep and goat fibroblasts *in vitro*. Nine of them showed significantly higher luciferase transcriptional activation compared to the pGL3-Promoter control (t-test, *P* value < 0.05) (Fig. 2I). Collectively, these results suggest that the active-RSCNEs potentially serves as enhancers in the process of rumen development and evolution.

### Conserved Non-coding Element interpretation method by Gene Regulatory Network (CNEReg)

After demonstrating that active-RSNCEs may often function as enhancers and hence have significant impacts on morphological evolution [21], we next developed CNEReg as an evolutionary Conserved Non-coding Element interpretation method. The method works by modeling the paired gene expression and chromatin accessibility data during rumen and esophagus development and consolidating them into a GRN. A GRN helps to understand in detail the process of TF binding to active-RSCNEs, and how this leads to the cell type specific activation of RSEGs during different stages of development. CNEReg takes as input a set of paired time-series gene expression and chromatin accessibility data, ruminant comparative genomes, and comparative transcriptomes, and outputs the projected developmental regulatory network of the active-RSCNEs. The three major steps of CNEReg includes: multi-omics data integration, model component identification, and developmental regulatory network inference (Fig. 3A; Methods). The developmental regulatory network reconstruction is illustrated in the following sections.

**Fig. 3.**
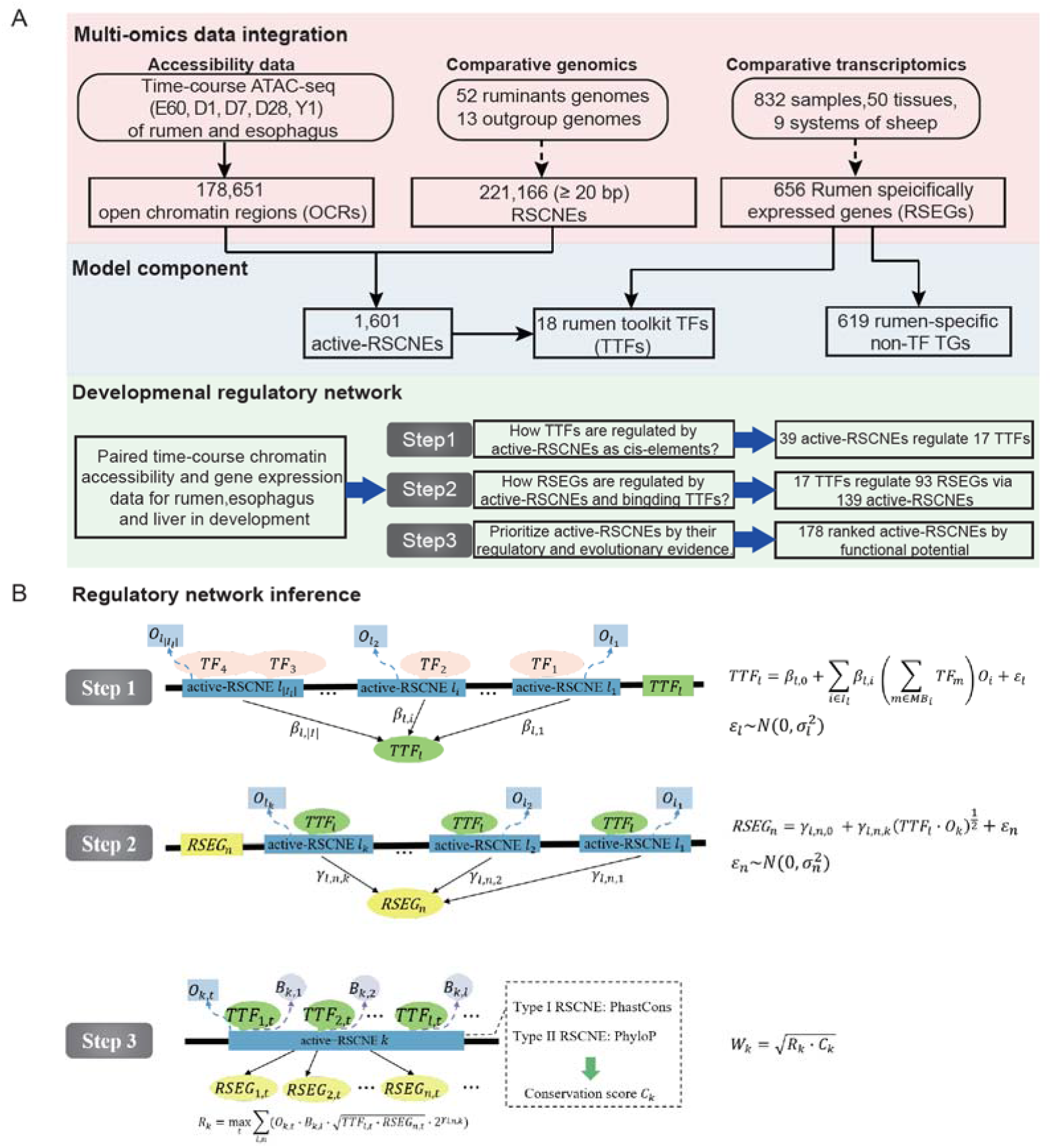
CNEReg interprets RSCNEs by reconstructing developmental regulatory network. (**A**) CNEReg inputs paired time-series gene expression & chromatin accessibility data, ruminant comparative genomes, and comparative transcriptomes and outputs the developmental regulatory network of active-RSCNEs. Three major steps of CNEReg includes: multi-omics data integration, model component identification, and developmental regulatory network inference. (**B**) The developmental regulatory network reconstruction is further illustrated in three steps. Step1: inferring the upstream regulations of rumen toolkit TFs (TTF). Step2: inferring the TTF’s downstream regulation to target genes via active-RSCNEs. Step3: Deriving active-RSCNE’s functional influence score by integrating regulatory strength in network and evolutionary conservation score. CNEReg’s model component and notations are detailed in Table 1.

**Table 1.** CNEReg model component and notations.

| Data and variables | Notation | Example |
| --- | --- | --- |
| Expression of TTF | $TTF_{l,t} :=$ expression of the $l$-th TTF on $t$-th time point | $TTF_{HOXC8} = 25.48$ on D7 in rumen |
| Expression of TF | $TF_m :=$ expression of the $m$-th TF | $TF_{JUN} = 1035.79$ on D7 in rumen |
| Expression of RSEG | $RSEG_{n,t} :=$ expression of the $n$-th RSEG on $t$-th time point | $RSEG_{SLC14A1} = 42.34$ on D7 in rumen |
| Accessibility of active-RSCNE | $O_{k,t} :=$ openness of $k$-th active-RSCNE on $t$-th time point | $O_{Chr1:196579342-242} = 18.83$ on D7 in rumen |
| TFs with motif match in an active-RSCNE | $MB_i :=$ set of TFs with significant motif match in $i$-th active-RSCNE | HOXC8 has motif match at active-RSCNE<br>$Chr1:196579342 - 242$ |
| Motif matching strength of TF on RE | $B_{i,l} :=$ sum of $-\log(p\text{-value})$ of $l$-th TF's motif on $i$-th active-RSCNE | $B_{Chr1:196579342-242} = 4.28486$ |

### Identifying toolkit transcription factors

We proposed toolkit transcription factors (TTFs) as the core concept of CNEReg and developed a computational pipeline to discover the developmental genetic toolkit TFs in evo-devo which may controls development, pattern formulation, and identity of body parts (details in Methods). We first separated 37 TFs from 619 non-TF target genes (TGs) in 656 RSEGs. Those 37 TFs are further filtered by a more stringent expression specificity *JMS* score and are required to have nearby active-RSCNEs in the upstream or downstream 1 Mbp to TSS (Methods). Finally, 18 TTFs are defined (Additional file 1: Table S5) and their expression profile phylogeny well recovers the tissue lineages system (Fig. 4A). Rumen was clustered the closest to reticulum, omasum, and esophagus and then skin and other keratin tissues, which is consistent with the basic stratified epithelium shared in rumen with skin. These 18 TTFs also well represented rumen’s major functions associated with other tissue systems, including gastrointestinal system, integumentary system, reproductive system, muscular system, nervous system, and endocrine system (Fig. 4B).

**Fig. 4.**
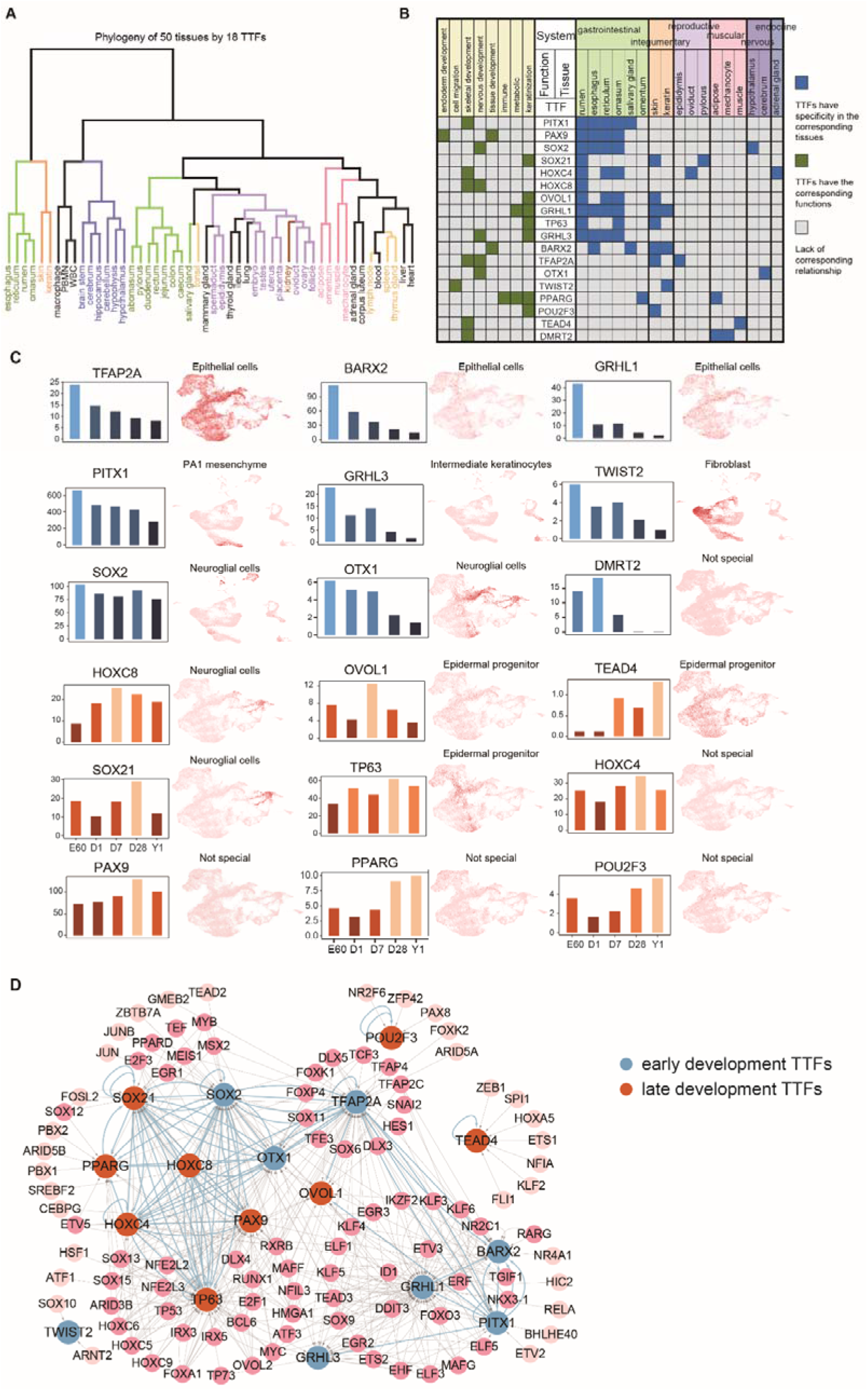
18 rumen TTFs and its upstream regulations. (**A**) Phylogeny of 50 tissues from sheep by 18 rumen TTFs’ expression groups the samples well by different lineages and biological system. (**B**) 18 rumen TTFs’ biological functions (marked by green) and the tissue with high expression (marked in blue). Tissues are grouped and colored by their lineages. (**C**) 18 rumen TTFs’ expression values along the development stages. By their dynamic patterns, they can be grouped into early (cold colored) and late (warm colored). In addition, 18 rumen TTFs’ expression in skin organoids scRNA-seq data are visualized by Uniform Manifold Approximation and Projection (UMAP) plot. The associated specific cell type names are labeled. (**D**) Rumen TTFs’ upstream gene regulatory network shows the candidate TFs with statistical significance. Nodes are colored by early and late TTFs. Blue edges highlight the regulatory relationship among TTFs.

We observed that rumen recruited TTFs from multiple tissues to drive gene expression and expressed more TTFs from gastrointestinal system than other systems. For example, paired box protein 9 (PAX9) is a known key transcription factor during esophagus differentiation, which may play an important role in rumen’s origin from the esophagus [22]. The homeobox family TFs HOXC8 and HOXC4, together with PITX1 are key developmental regulator for specific positional identities on the anterior-posterior axis [23, 24]. The other four TTFs, OVOL1, SOX21, TFAP2A, TP63 are from integumentary system and serve as master regulators in the regulation of epithelial development and differentiation [25–28].

We classified 18 TTFs into two types according to their dynamic gene expression pattern during rumen development. PITX1, BARX2, SOX2, GRHL1, GRHL3, TFAP2A, OTX1, DMRT2, and TWIST2 are early development TTFs showing the highest expression at E60 or D1 (Fig. 4C). In contrast, PAX9, TP63, HOXC4, SOX21, HOXC8, OVOL1, PPARG, POU2F3, and TEAD4 are late development TTFs and highly expressed at D7, D28, or Y1 (Fig. 4C). We further associated those TTFs with 6 cell types by their expression level in skin organoids scRNA-seq data [29]. The organoid culture system presents a complex skin organ model by reprogramming pluripotent stem cells. For example, TFAP2A is specifically expressed in epithelial cells (Fig. 4C).

### Constructing TTFs’ upstream and downstream regulations

To explore how TTFs are regulated and recruited, we scanned the active-RSCNEs near TTFs for the sequence-specific TF’s motif binding, retained those TFs correlating well with TTFs (Spearman’s correlation coefficient > 0.6 across RNA-seq samples), and fitted a linear regression model integrating our paired expression and chromatin accessibility data to reveal 18 TTFs’ upstream regulators (Fig 3B; Methods). The resulting TTFs’ upstream regulatory network (Fig. 4D) identified 39 active-RSCNEs (15 Type I and 24 Type II) bound by 113TFs for 18 TTFs (Additional file 1: Table S6). GRHL1, an important regulator in keratin expression [30], is regulated by 31 TFs via 6 active-RSCNEs, suggesting its potential roles in rumen development.

To explore 18 TTFs’ regulatory roles, we first scanned 1,440 active-RSCNEs located 1 Mbp upstream or downstream around 512 RSEGs (FPKM > 1 in at least one development stage) by HOMER [31] for binding sites of the 18 rumen TTFs. Then linear regression model quantitatively associated the accessibility of active-RSCNEs with the expression of TTFs and RSEGs (Fig. 3B; Methods). The resulting TTFs’ downstream regulatory network linked 139 active-RSCNEs (26 Type I and 113 Type II) with 17 TTFs and 93 RSEGs (Fig. 5A; Additional file 1: Table S7). RSEGs were categorized into different tissue systems. The gastrointestinal and integumentary systems both have 28 RSEGs which are functionally enriched in hair/molting cycle process (Fisher’s exact test, adjusted-*P* value = 1.50×10^−2^) and regulation of antimicrobial peptide production (Fisher’s exact test, adjusted-*P* value = 3.58×10^−6^). This is consistent with our previous finding that rumen evolved several important antibacterial functions specifically managing the microbiome composition [2]. *SLC14A1* gene was specifically highly expressed in the rumen and hypothesized to be recruited from the urinary system (Fig. 5A). CNEReg identified four active-RSCNEs bound by three TTFs, OTX1, PPARG, and SOX21, to regulate *SLC14A1* (Fig. 5B).

**Fig. 5.**
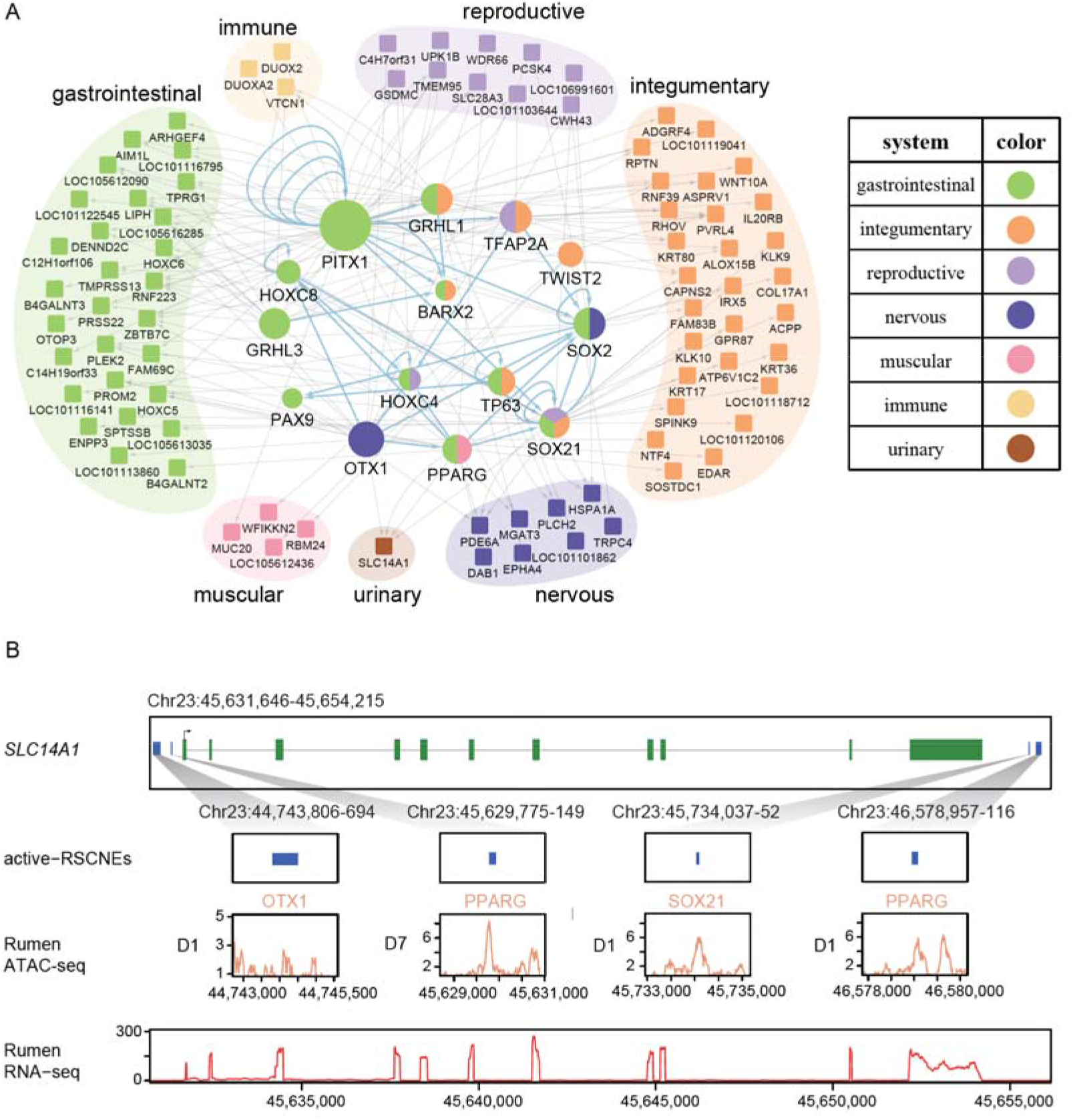
Rumen TTFs’ downstream regulatory network. (**A**) Rumen TTFs’ downstream regulatory network with 17 rumen TTFs regulating 93 TGs via 139 active-RSCNEs. TTFs are colored by the tissue they are highly expressed and TGs are annotated and colored by their biological system. (**B**) An example from the regulatory network shows that *SLC14A1* is regulated by four active-RSCNEs with TTFs’ motif occurrence. The expression and chromatin accessibility tracks are derived from rumen ATAC-seq (D1 or D7) and RNA-seq data (Y1).

CNEReg designed a functional influence score by integrating regulation and conservation in evolution (Fig 3B; Methods) and ranked the active-RSCNEs in TTF’s upstream and downstream networks (Additional file 2: Fig. S4, 5; Additional file 1: Tables S6, 7). Then we selected top 10 active-RSCNEs for enhancer activity detection in sheep fibroblasts in *vitro*. 9 of 10 showed significantly higher luciferase transcriptional activation compared to the pGL3-Promoter control (t-test, *P* value < 0.05) (Additional file 2: Fig. S6). Collectively, CNEReg provides high quality developmental regulatory network to study rumen evolution.

### Regulatory sub-network underlying rumen and the esophagus divergence

We previously hypothesized that the anatomical predecessor of the rumen is the esophagus based on their similar expression profile compared to other 49 tissues [5, 7]. It is therefore of interest to identify the gene regulatory network underlying the differentiation between rumen and esophagus. We first identified differentially expressed genes (4, 258, 577 and 2,372, for E60, D1, D7, and Y1 in Additional file 2: Fig. S7A) and differentially accessible regions (9,436, 10,004, 3,984, 3,566 and 26 for E60, D1, D7, D28, and Y1 in Additional file 2: Fig. S7B) between rumen and esophagus at each developmental stage. Then, we identified six TTFs (PPARG, SOX21, TP63, OTX1, SOX2, and HOXC8) showing both significant differences in expression and in motifs enriched within the rumen OCRs (Fig. 6A; Methods). HOXC8 shows the largest difference at the earliest developmental stage, both in expression level and motif enrichment, and SOX21, SOX2, OTX1 and PPARG show similar trends. TP63 differentiates from D7 where the gene expression level and motif enrichment decline quickly in esophagus but not in rumen.

**Fig. 6.**
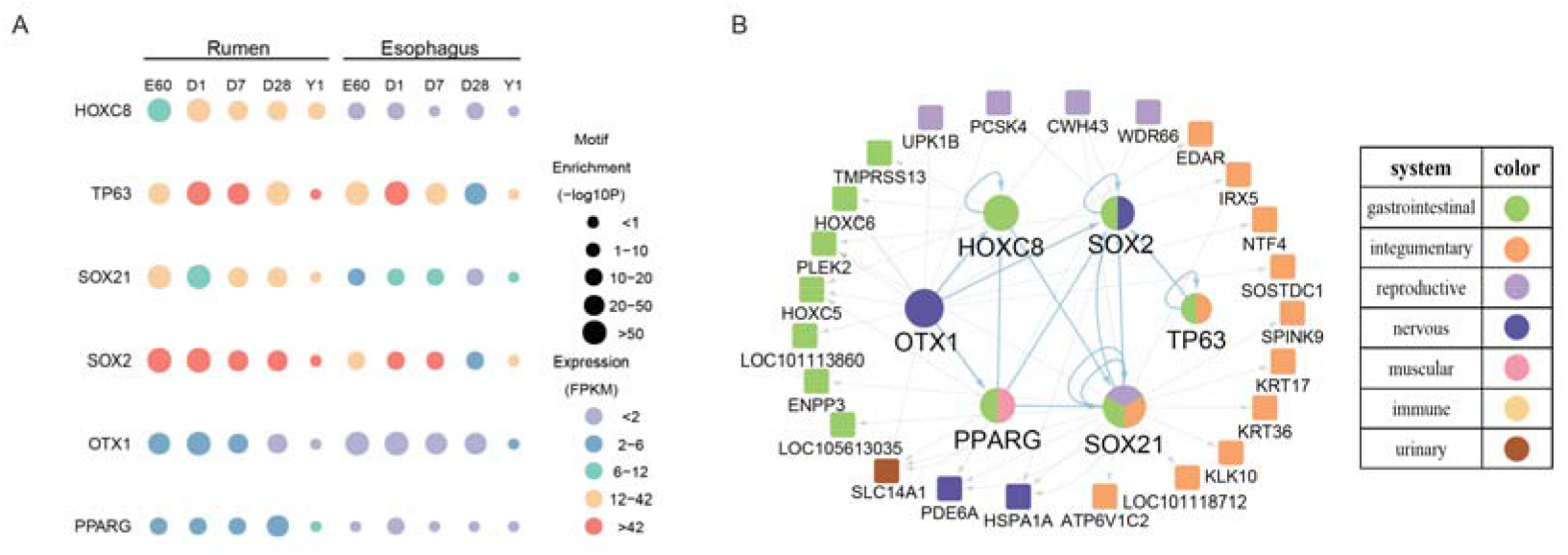
Regulatory network sheds lights on the difference between rumen and esophagus in development. **(A)** Dynamics across stages for the 6 differential TTFs between rumen and esophagus by integrating motif enrichment in differential ATAC-seq peaks and gene expression level. **(B)** 6 differential rumen TTFs’ downstream regulatory subnetwork, which hypothesizes that rumen evolves from homologous tissue esophagus by functional innovation through recruiting OTX1, SOX21, HOXC8, SOX2, TP63, PPARG and utilizing 16 active-RSCNEs to rewire developmental regulations.

We extracted the six differential TTFs from the TTFs downstream regulatory network to form a regulatory sub-network which also including 24 differentially expressed RSEGs and 38 active-RSCNEs (Fig. 6B; Additional file 1: Table S8). The 24 differentially expressed RSEGs were classified into gastrointestinal, integumentary, reproductive, nervous, muscular, immune, and urinary systems and 10 of 24 non-TF RSEGs were classified into integumentary system. Seven non-TF RSEGs (*KRT17*, *KRT36, LOC101118712, ATP6V1C2*, *KLK10, SPINK9*, and *IRX*) were regulated by SOX21. Previous study revealed that SOX21 could determine the fate of ectodermal organ and control the epithelial differentiation [28]. We observed that SOX21 binds to “Chr11:40325877-150” to regulate the expression of KRT17, KRT36 and LOC101118712. The functional influence of “Chr11:40325877-150” is ranked at the top of all Type II active-RSCNEs in the differentially regulatory sub-network (Additional file 1: Table S8). Those RSEGs were enriched in epidermis development, formation of anatomical boundary, and urea transmembrane transport biological process (Additional file 1: Table S9), which are consistent with the function difference between rumen and esophagus. The 38 active-RSCNEs may imply the potential genetic basis of rumen’s origin and evolution from esophagus.

### Transposable element may rewire gene regulatory network through active-RSCNEs

After interpreting active-RSCNEs as important regulators for TTFs and RSEGs in rumen development, we next address the genomic origin of the active-RSCNEs. Transposable elements (TE) are known to constitute a high proportion of taxonomy-specific CNEs, play a central role in rewiring gene regulatory networks, and to facilitate novel or rapid evolution of ecologically relevant traits [32, 33]. Hence, we estimated which active-RSCNEs may derive from TEs. Among 39 and 139 active-RSCNEs in TTF’s upstream and downstream networks, we identified six (15.38%) and 12 (8.6%) TEs, respectively. This gives a 1.8-fold enrichment of TEs in active-RSCNEs associated with TTFs relative to non-TTF RSEGs. At the gene level, six of 18 TTFs (33.33%) and 12 of 93 RSEGs (12.90%) are regulated by TE via active-RSCNEs. This gives a 2.58-fold enrichment. As a background, there are 85 TEs around all 656 RSEGs (+/− 200 kbp) and give an average 13%. Together, our data suggests that TE may recruit expression of TTFs and rewiring the regulatory network to give rise of trait novelties.

## Discussion

The evolution of new trait is driven by several types of genetic reprogramming, including mutations in protein-coding genes and post-transcriptional mechanisms, transformation of regulatory elements such as promoters and enhancers, and recruitment of gene expression from other organs [34, 35]. Mutations in non-coding regulatory regions are believed to selectively perturb target gene expression in specific tissue context and thereby circumvent any pleiotropic effects from protein-coding mutations [36]. Recent advances in comparative genomics, along with the increased availability of whole genome sequences, have led to the identification of many conserved non-coding elements (CNEs), which are assumed to have regulatory functions [1, 6, 37]. Therefore, the time is ripe for an analytical framework to investigate the regulatory role of such CNEs.

Here we propose a model of gene expression recruitment by CNEs. Our results show how CNEs can regulate gene expression as either *trans*-regulatory elements (TTF in our study) or *cis*-regulatory elements (active-RSCNEs) of target genes (RSEGs). CNEReg provides a framework to integrate comparative genomics, comparative transcriptomic, and multi-omics data to interpret CNEs by GRN. On one hand, GRN presents the global picture how rumen recruits gene expression from other tissues by activating RSCNEs to achieve many traits. On the other hand, GRN identifies TTFs and active-RSCNEs as hypotheses, which need to be pursuit by in *vitro* and in *vivo* functional studies. Our method for systematically interpreting conserved *cis*-regulatory sequence in non-coding region by integrating developmental multi-omics data will have broad interest in other applications. For example, the Zoonomia Project describes a whole-genome alignment of 240 species comprising representatives from more than 80% of mammalian families [38]. The Bird 10,000 Genomes (B10K) Project provides comparative genome dataset for 363 genomes from 92.4% of bird families [39]. Recently ~6.9 million CNEs from many vertebrate genomes are collected into dbCNS and await to be interpreted [40].

Our work is limited in several aspects. CNEReg infers the gene regulation as the interaction of TFs with accessible DNA regions in development and relies on the correlation of gene expression and chromatin accessibility across samples. Much deeper understanding can be revealed by ChIP-seq data and 3D chromatin interaction data to provide physical enhancer promoter interactions. In addition, time course regulatory analysis on the omics data measured at shorter and closer developmental stages will help [12]. Furthermore, developmental samples are known as a heterogeneous mixture of many cell types and it will be fruitful to infer the GRNs of the underlying cell types based on scATAC-seq and scRNA-seq data [10].

## Conclusions

In conclusion, CNEReg is demonstrated as a systematic approach to understand the large-scale maps of CNEs by modeling omics data over development for its act on gene regulation. We see the potential that CNEReg can be generalized to understand the complex traits or the origin and evolution of vertebrate organs with multi-omics data generated in proper time and space. Our method allows evo-devo thinking in how gene regulation could evolve and shape animal evolution.

## Methods

### CNEReg infers developmental regulatory network to interpret conserved non-coding element

CNEReg aims to systematically fill the gap between conserved non-coding elements (CNEs) and its significantly impacted morphology in evolution. This is done by reconstructing a developmental regulatory network by paired time series of paired gene expression and chromatin accessibility data. Particularly in sheep CNEs are RSCNEs and morphology is the innovation of rumen organ, which is further denoted by the set of rumen specific genes RSEGs. We reconstruct gene regulatory network during rumen development to systematically understand how the TFs regulate genes via batteries of RSCNEs, which over development, lead to the cell type specific activation of RSEGs.

The main idea of CNEReg is to focus on those toolkit TFs as major players in evo-devo to study how those TFs are regulated by RSCNEs and how they utilize RSCNEs to regulate RSEGs. CNEReg models the expression of target genes (TG) conditional on chromatin accessibility of RSCNEs and expression of transcription factors (TF). CNEReg is composed by three steps as shown in Fig. 3 and uses three formulations to model, (1) expression of toolkit TFs, (2) expression of RSEGs, (3) functional influence of RSCNEs (Fig. 3; Table 1).

### Step 1. Modeling expression of toolkit transcription factors (TTFs)

We first identify toolkit TFs by its nearby evolutionally conserved cis-regulatory element in genome, expression pattern across tissues, expression levels in developmental stages. TTFs should satisfy four conditions: (1) TFs should be rumen specifically expressed genes (37 TFs in the 656 RSEGs), (2) there should be active-RSCNEs around TFs (+/− 1M bp, 35 TFs remains), (3) TFs should be expressed (FPKM > 1) in at least one time point during rumen development (30 TFs remains), and (4) these TFs should be additional tissue specificity. TFs were ranked by our tissue specificity score *JMS* and only the TFs for top 50 specificity in at least one tissue will be selected (18 TFs remains). Finally, 18 TFs were identified as TTFs and listed in **Supplementary text**). These TFs played a leading role in rumen development (Additional file 1: Table S5) and served as the main component to construct the rumen developmental regulatory network.

Next, we model how the TTFs are regulated from paired gene expression and chromatin accessibility data, i.e., to reconstruct the upstream regulatory network of TTFs. We established a linear regression model as follows to explore the upstream regulatory network of the 18 TTFs (Schematic illustration in **Fig. 3** and mathematical notations in **Table 1**).

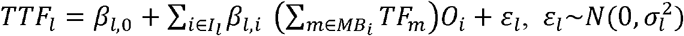

where *TTF_l_* is the expression of the *l*-th TTF; *MB_i_* is the set of TFs with significant motif match in the *i*-th active-RSCNE; *TF_m_* is the expression of the *m*-th candidate TF with binding motif to regulate the *l*-th TTF. The Spearman correlation coefficient between *TF_m_* and *TTF_l_* is greater than 0.6 (FDR *q*-value < 0.01) to ensure the potential regulatory relationship; *O_i_* represents the chromatin accessibility score of the *i*-th active-RSCNE within 2 Mbps around the *l*-th TTF. *β* is the parameter to be estimated. If *β_l,i_* is statistically significant in the regression analysis, the *i*-th active-RSCNE and its TFs in *MB_i_* will be contained in the upstream regulatory network of the *l*-th TTF.

### Step 2. Modeling expression of rumen specifically expressed genes (RSEGs)

We model how the RSEGs are regulated by TTFs and its active-RSCNE from paired gene expression and chromatin accessibility data, i.e., to reconstruct the downstream network regulated by TTFs. We established the linear regression model as follows (Schematic illustration in **Fig. 3** and mathematical notations in **Table 1**).

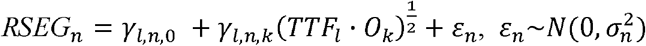

where *TTF_l_* is the expression of the *l*-th TTF; *O_k_* represents the chromatin accessibility score of the *k*-th active-RSCNE with binding sites of the *l*-th TTF; *RSEG_n_* is the expression of the *n*-th *RSEG* with the *k*-th active-RSCNE around within 2 Mbps. In practice, we determine the downstream regulation relationship with Spearman correlation that can eliminate the outlier values to simplify the calculation. When the Spearman correlation coefficient *γ_l,n,k_* between *RSEG_n_* and (*TTF_l_ · Ok)12* is greater than 0.7 (FDR *q*-value < 0.01), the *n*-th RSEG is likely to be regulated by the *l*-th TTF through binding on the *k*-th active-RSCNE. The extracted TTF, active-RSCNEs, and RSEGs triplets are the TTF’s downstream regulatory network.

### Step 3. Quantifying functional influence of active-RSCNEs

We finally quantify the functional influence of active-RSCNEs, rank the active-RSCNEs, and select the top active-RSCNEs as experimental candidates. This task can be done by integrate the RSCNE’s conservation score in evolution with its regulatory potential in the developmental regulatory network.

We firstly collected conservation scores of active-RSCNEs from comparative genomics study [5]. RSCNEs were classified into two types by their conservation patters across species. Type I RSCNEs had no outgroup sequence aligned and Type II RSCNEs had orthologous sequences in one or more outgroups but were only conserved in ruminant. For the *k*-th active-RSCNE, the conservation score *C_k_* was calculated by PhastCons score (Type I) or PhyloP score (Type II).

We then estimated the regulatory strength of active-RSCNEs in the upstream and downstream regulatory network of TTFs. An active-RSCNE played a regulatory role in the regulatory network if four conditions were satisfied: (1) this active-RSCNE should be a chromatin accessible peak, (2) TTFs should bind on this active-RSCNE, (3) RSEGs regulated by this active-RSCNE with TTFs binding should be expressed, and (4) the expression of binding TTFs and the accessibility of this active-RSCNE should be correlated with the expression of regulated RSEGs. By combining these four factors, we defined the regulatory strength *R_k,t_* of the *k*-th active-RSCNE at time point *t* in the regulatory network as follows:

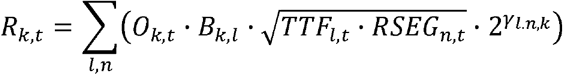

Where, *O_k,t_* is the chromatin accessibility score of the *k*-th active-RSCNE at time point *t* in rumen; *B_k,l_* is the motif binding strength of the *l*-th TTF on the *k*-th active-RSCNE (computed by HOMER); *TTF_l,t_* is the expression of the *l*-th TTF at time point *t* in rumen; *RSEG_n,t_* is the expression of the *n*-th RSEG at time point *t* in rumen; *γ_l,n,k_* is the Spearman correlation coefficient between *RSEG_n_* and 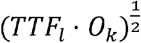 from the regulatory network. Then the regulatory strength *R_k_* of the *k*-th active-RSCNE was defined as the maximum value across all time points in rumen samples:

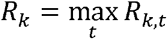

The regulatory strength *R_k_* is from the multi-omics data in development and conservation score *C_k_* is from multi-genome data across species. The two measures are respectively at regulation level and genome sequence level and can be naturally assumed independent to each other. In addition, we found that the regulatory strength and the conservation score were quite complementary to each other (Additional file 2: Fig. S4, 5) for active-RSCNEs. Hence, we defined the functional influence *W_k_* of the *k*-th active-RSCNE as the geometric mean of the regulatory strength *R_k_* and the conservation score *C_k_* as follows:

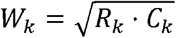

This functional influence score allows us to prioritize active-RSCNEs by importance in rumen innovation.

### Hierarchical clustering and principal component analysis (PCA)

We performed hierarchical clustering on the gene expression and peak chromatin accessibility profiles in 14 rumen samples at five time points (E60/D1/D7/D28/Y1). Heatmap was plotted by R package “pheatmap” with “correlation” as distance measure and “complete” as clustering method. Then we performed dimensional reduction by principal component analysis (PCA) with R function “prcomp”. The gene expression and chromatin accessibility value were log transformed as log_2_ (FPKM + 1) and log_2_ (Openness + 1) as input. FPKM is the reads per kilobase per million mapped reads and openness score was calculated for each peak under each condition as the fold change of reads number per base pair [10]. The first two principal components are shown in **Fig. 1D and E**.

### Definition of tissue specificity score

Specificity illustrates the property that gene are functional in one particular biological context compared to other contexts. For our transcriptomics data across 50 tissues in sheep, genes highly expressed in only one or several tissues but not expressed in other tissues were defined as tissue specific. Our gene expression matrix is with 23,126 rows (the number of expressed genes) and 830 columns (the number of samples sequenced in 50 sheep tissues, and each tissue has several biological replicates) (Additional file 1: Table S10).

To quantify the tissue specificity, we proposed a JMS score for a gene in certain tissue to combine gene expression level with a Jensen–Shannon Divergence (*JSD*) value as follows.

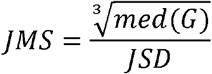

where *med*(*G*) represents gene’s median expression in a certain tissue. 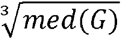 can guarantee that the numerator and denominator are on the same magnitude. *JSD* is the Jensen–Shannon divergence to evaluate the gene’s expression specificity introduced in [41]. It adopts an entropy-based measure to assess the similarity between two probability distributions in statistics as follows,

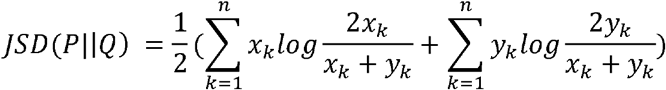

Where *P* = (*x*_1_, *x*_2_, … *x_n_*) and *Q* = (*y*_1_, *y*_2_, … *y_n_*) are two probability distributions constructed from our gene expression values across tissues. n is the number of samples. Given each row of our gene expression matrix, we then normalized the gene’s expression vector, i.e., each element in this vector was divided by the sum of all elements. For a given gene, *Q* = (*y*_1_, *y*_2_, … *y_n_*) is its corresponding normalized row vector. Given the tissue we are interested, *P* = (*x*_1_, *x*_2_,… *x_n_*) is constructed as a control vector whose components are 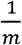 in the given tissue with *m* replicates and 0 in other tissues. Finally, the *JSD* will be calculated as the divergence between P and Q for a certain gene in certain tissue. The smaller the *JSD* value, the more specific this gene in this tissue.

In summary, *JSD* provided a relative specificity score by a nonlinear measure of divergence. We further extended it by emphasizing significantly highly expressed genes in certain tissues to enhancing specificity. This JMS score allows us to better explore the toolkit transcription factors’ expression patterns and recruitment of genes based on tissue specificity.

### Differential regulatory network construction between rumen and esophagus

We constructed differential regulatory network between rumen and esophagus by extracting differential RSEGs, differential TTFs, and active-RSCNEs associated sub-network from the regulatory network of TTFs. The differential RSEGs and differential TTFs are defined as follows.

#### Differential RSEGs between rumen and esophagus

We used R packages “limma” and “edgeR” to extract differential genes at four developmental time points (E60/D1/D7/D28) with thresholds FDR < 0.05 and log_2_FC > 1 (FC was fold-change of FPKM in rumen relative to esophagus). It was noted that at time point Y1, we only had one biological replicate for RNA-seq data in rumen and esophagus separately and we could not perform F-test on these two samples. Instead, we identified genes with FPKM > 2 in rumen and FC > 2 as differential genes. Then we combined differential genes at five time points to get differential genes set between rumen and esophagus. Differential RSEGs between rumen and esophagus were intersection of differential genes set and RSEGs set in regulatory network of TTFs.

#### Differentially accessible peaks between rumen and esophagus

We implemented R packages “limma” and “edgeR” to get differentially accessible peaks between rumen and esophagus at five developmental time points (E60/D1/D7/D28/Y1) with thresholds FDR < 0.05 and |log_2_FC| > 1 (FC was fold-change of chromatin accessibility score in rumen relative to esophagus).

#### Differential TTFs between rumen and esophagus

We first collected 1,027 TFs of sheep from animalTFDB3.0 (http://bioinfo.life.hust.edu.cn/AnimalTFDB/#!/). The 15,835 expressed genes in rumen and esophagus were intersected with these 1,027 TFs to obtain 768 TFs for the following analysis. We used HOMER to find TFs’ binding on the differentially accessible peaks with threshold −log_10_*p*value > 6 at each time point. Then we used R packages “limma” and “edgeR” to get differentially expressed TFs at four time points (E60/D1/D7/D28) with threshold FDR < 0.05 and log_2_FC > 1. We identified differentially expressed TFs at time point Y1 with threshold FPKM > 2 in rumen and FC > 2. Differential TFs set was defined as the intersection of TFs binding on differentially accessible peaks and differentially expressed TFs. Differential TTFs between rumen and esophagus were intersection of differential TFs set and TTFs set in regulatory network of TTFs.

### Collecting samples for ATAC-seq and RNA-seq

We collected a total of 37 samples of the rumen, esophagus epithelium tissues and liver tissues from 14 Hu sheep including 5 time points (embryo 60-day, 1-day, 7-day, 28-day, and 1-year) from XiLaiYuan ecological agriculture co. LTD in Taizhou city (Jiangsu, China). All samples rinsed with PBS and were soaked in cold 1×PBS added with penicillin-streptomycin. All animals were slaughtered under the guidelines of Northwest A&F University Animal Care Committee.

### ATAC-seq library preparation, sequencing, and analysis

#### Isolation of ruminal and esophageal epithelial cells

A piece of ruminal epithelial tissue was removed from PBS buffer (pH 7.4), placed on a watch glass, and brushed with sterile D-Hanks in all directions. The clipped tissue (approximately 500 mg) was placed in a small beaker and rinsed 2-3 times with a D-Hanks (pH 7.4) solution (4 times antibody, pre-warmed in a 37 °C water bath). Next, 0.25% trypsin (15 ml) was pre-warmed in a 37 °C water bath and added to a conical flask with the rumen epithelium sample, which was then digested in a 37 °C water bath for 30 min while shaking well every 5 min. The ruminal epithelial tissue was removed and the trypsin digestion solution was discarded. This step was repeated three times until the epithelium felt sticky. The treated epithelial tissue was then placed in a sterile beaker and rinsed three times with D-Hanks solution, and this step was repeated using a fresh beaker. Ten milliliters of trypsin were added, and the mixture was digested in a 37 °C water bath for 10-20 min until the cells detached. Cells from the first 3-4 detachments were not collected because these are generally necrotic or granular cells. Only the last two digested cell types (spinous and basal cells) were generally collected, after the cells in the digested sample were observed under a microscope. The cells were filtered through a cell strainer and added to a 10 ml centrifuge tube containing a drop of calf serum. The above digestion and collection steps were repeated 3 times. The digestate was collected following centrifugation at 1500 r/min for 5-10 min, and the supernatant was discarded. One milliliter of Dulbecco’s Modified Eagle Medium (DMEM) solution was added to the precipitate, and the mixture shaken or blown to adjust the cell density to 10^6^ cells/ml. Trypan blue was added to verify that cell viability reached 95%. The esophageal epithelial cells were obtained with the same pipeline with the ruminal epithelial cells as above.

#### Preparation of nuclei

To prepare nuclei, we spun 50,000 cells at 500 x*g* for 5 min and then washed the pellet using 50 μl of cold 1× PBS. The solution was then centrifuged at 500 x*g* for 5 min, and the cells were lysed using cold lysis buffer (10 mM Tris-HCI, pH 7.4, 10 mM NaCl, 3 mM MgCl2 and 0.1 NP40). Immediately after lysis, the nuclei were spun at 500 x*g* for 10 min using a refrigerated centrifuge. To avoid losing cells during the nucleus preparation, we used a fixed angle centrifuge and carefully pipetted away from the pellet after centrifugation.

#### Transposition and purification

The pellet was immediately resuspended in transposase reaction mix (17.5 μl of DEPC H_2_O, 5μl of TTBL buffer, 2.5 μl of TTE mix buffer and all the nuclear DNA). The transposition reaction was carried out for 10 min at 55 □ in metal bath, and the sample was immediately purified using a Qiagen MinElute kit.

#### Library construction

PCR was performed to amplify the library for 14 cycles using the following PCR conditions: 72 °C for 3 min, 98 °C for 30 s, and thermocycling at 98 °C for 15 s, 60 °C for 30 s and 72 °C for 3 min.

#### Data quality control and short-read alignment

Sequencing reads must undergo quality control and adapter trimming to optimize the alignment process. FastQC (version 0.11.5) [42] was used to assess overall quality. Reads were trimmed for quality as well as the presence of adapter sequences using the Trim Galore Wrapper script [43] with default parameters. Raw ATAC-seq reads of sheep were mapped to the sheep reference genome (NCBI assembly Oar_v4.0) using Bowtie2 (version 2.2.8) [44] with default parameters. Duplicated reads were removed using the default parameters in Picard (version 2.1.1). Reads mapping to mitochondrial DNA were excluded from the analysis together with low-quality reads (MAPQ < 20).

#### Open accessible peak calling

Accessible regions and peaks were identified using MACS [45] with parameters “-q 0.05 -shift 37 -extsize 73” for narrow peaks. The centers of identified peaks were used to define peak overlaps with genomic features according to the following criteria. If a center site was located in the promoter of a gene (2 kbp upstream from the transcription start site (TSS)), or the gene body, the peaks would be assigned to that gene. Distal intergenic regions refer to regions > 3 kbp from the TSS and > 1 kbp from the transcription termination site (TTS).

#### Consensus peaks analysis

Open accessible peaks were identified in four biological replicates of each tissue by using “bedtools intersect”, and consensus peaks with openness values of each peak in each sample were built by merging these regions and calculated with R package “Diffbind” (version 2.10.0) [46].

#### Peak annotation

Peak annotation was performed using R packages “GenomicFeatures”, “ChIPseeker”, and “AnnotationHub”.

### RNA-seq library preparation and sequencing

We prepared directional RNA-seq libraries from the cells of the same samples as used for ATAC-seq. Each sample was added 1ml Trizol protocol (Invitrogen, USA), and frozen in −80 °C until utilization.

#### RNA isolation, library construction, and sequencing

In all tissue samples collected for this study, total RNA was isolated from a frozen sample according to the Trizol protocol (Invitrogen, USA), using 1.5 μg RNA per sample as the input material for sample preparation. Sequencing libraries were generated using a NEBNext® Ultra RNA Library Prep Kit for Illumina® (NEB, USA) according to the manufacturer’s recommendations, and index codes were added to attribute sequences to samples. Briefly, mRNA was purified from total RNA using poly-T oligo-attached magnetic beads and fragmented using divalent cations at elevated temperature in NEB Next First-Strand Synthesis Reaction Buffer (5X). First-strand cDNA was synthesized using random hexamer primers and M-MuLV Reverse Transcriptase (RNase H). Second-strand cDNA was subsequently synthesized using DNA Polymerase I and RNase H. Remaining overhangs were converted into blunt ends by exonuclease/polymerase activity. After adenylation of 3’ ends of DNA fragments, NEB Next Adaptors with hairpin loop structures were ligated to prepare for hybridization. To select cDNA fragments with appropriate lengths, the library fragments were purified with an AMPure XP system (Beckman Coulter, Beverly, USA). Then 3 μl of USER Enzyme buffer (NEB, USA) was incubated with size-selected, adaptor-ligated cDNA at 37 °C for 15 min followed by 5 min at 95 °C before PCR amplification, using Phusion High-Fidelity DNA polymerase, Universal PCR primers, and Index (X) Primer. Finally, PCR products were purified using the AMPure XP system, and library quality was assessed using an Agilent Bioanalyzer 2100 system. The index-coded samples were clustered with a cBot Cluster Generation System using a HiSeq 4000 PE Cluster Kit (Illumina) according to the manufacturer’s instructions. After cluster generation, the library preparations were sequenced on an Illumina Hiseq X Ten platform, and 150 bp paired-end reads were generated. All these sequencing procedures were performed by Novogene Technology Co., Ltd., Beijing, China.

#### RNA-seq data quality control and quantification processing

We obtained high-quality reads by removing adaptor sequences and filtering low-quality reads from raw reads using Trimmomatic (version 0.36) [47] with the following parameters: LEADING:3 TRAILING:3 SLIDINGWINDOW:4:15 MINLEN:40. High-quality reads were all aligned to the NCBI assembly Oar_v4.0 reference sheep genome [48]. For this, we used STAR (Version 2.5.1) [49] with the following parameters: outFilterMultimapNmax 1, outFilterIntronMotifs RemoveNoncanonical Unannotated, outFilterMismatchNmax 10, outSAMstrandField intronMotif, outSJfilterReads Unique, outSAMtype BAM Unsorted, outReadsUnmapped Fastx, and outFileNamePrefix. The unmapped reads were extracted by SAMtools (Version 1.3) [50] for further mapping by HISAT2 (Version 2.0.3-beta) [51]. We computed Fragments Per Kilobase per Million mapped reads (FPKM) values for the genes in each sample using StringTie (Version1.3.4) [52].

As the samples were prepared and sequenced in three known distinct batches (see Additional file 1: Table S1), we used the *removeBatchEffect()* function from R *limma* package to build a linear model with the batch information and the cell types on log2-transformed FPKM+1, and we regressed out the batch variable.

#### Cell lines and cell culture

Conspecific cell lines can be used to validate the regulatory activity of RSCNEs. For example, mouse NHI3T3 fibroblast cells were used to validate the enhancer activity in mouse of one CNE which showed an ability of regulating the loss and re-emergence of legs in snakes [53]. Hence, we selected fibroblast cells of ruminants for *in vitro* regulatory activity experiments. Sheep and goat fibroblast cells were provided by Guangxi University and were cultured in Dulbecco’s Modified Eagle Medium (DMEM) containing 10% FBS (Gibco, Grand Island, NY, USA). All cell lines used in this study were maintained in the specified medium supplemented with 1 × Penicillin–Streptomycin (Gibco) and incubated in 5% CO2 at 37 °C.

#### Cloning and luciferase assays

All the reporter constructs were cloned into pGL-3 promoter plasmids (Promega, Madison, WI, USA). Fragments of the candidate RSCNEs were cloned into pGL3-promoter vector digested by *BamH* I and *Sal* □ downstream of the luciferase gene. All constructs were confirmed by sequencing. Transfection of all reporter plasmids constructs was performed using TurboFect (R0531, Thermo Scientific, Waltham, USA). Renilla Luciferase pRL-TK-Rluc (Promega) served as a transfection control, and luciferase expression was subsequently monitored with the dual luciferase assay (Promega) 24 h after transfection. Each luciferase assay was monitored at least five times, independently.

#### Statistics

The t-test in the GraphPad Prism7.0 software (Prism, San Diego, CA, USA) was applied to calculate the significance for the regulatory activity. Differences were statistically significant when *p* value < 0.05.

## Supplementary Information

**Additional file 1: Table S1.** Statistic of 37 ATAC-seq data used in this study. **Table S2.** Statistic of RNA-seq data used in our study. **Table S3.** 1,601 active-RSCNEs. **Table S4.** 1,061 active-RSCNEs are enriched for binding motifs of transcriptional regulators known to play vital role in rumen development (128 motifs with Benjamini q-value < 1.00×10−3). **Table S5.** 18 rumen toolkit TFs (TTFs). **Table S6.** Upstream regulatory network of 18 rumen toolkit TFs. **Table S7.** Downstream regulatory network of 18 rumen toolkit TFs. **Table S8.** Differential regulatory sub-network between rumen and esophagus. **Table S9.** GO enrichment analysis of 52 TGs in the differential regulatory sub-network between rumen and esophagus. BP denotes Biological Process, MF denotes Molecular Function, and CC denotes Cellular Component. **Table S10.** The gene expression profile of 655 rumen specifically expressed genes (RSEGs) which showed by FKPM value.

**Additional file 2: Fig. S1.** Data quality check for the ATAC-seq samples by their sequence depth, fragment distribution, and QC score. **Fig. S2**. Paired expression and chromatin accessibility time series data reveals the regulatory landscape for rumen and esophagus development. **Fig. S3.** Further characterization of active-RSCNEs. **Fig. S4.** Relationships between the regulatory strength and the conservation score of TTF upstream network. **Fig. S5.** Relationships between the regulatory strength and the conservation score of TTF downstream network. **Fig. S6.** Luciferase activity assays of 10 active-RSCNEs with top functional influence score. **Fig. S7.**Differentially expressed genes and differentially accessible peaks between rumen and esophagus at each stage.

## Declarations

### Acknowledgments

We thank High-Performance Computing (HPC) of Northwest A&F University (NWAFU) for providing computing resources.

### Authors’ contributions

Y.W, H.W, Y.J. and W.W. conceived the project and designed the research. X.P., Z.M, and X.S. performed the majority of analysis with contributions from Y.C., C.Z.; X.P. and T.Z. prepared rumen and esophagus epithelium cells and hepatocyte for ATAC-seq and RNA-seq. H.L. performed the luciferase reporter assay. X.P., Y.W., R.H., Z.M, and X.S. drafted the manuscript. All authors wrote, revised, and contributed to the final manuscript.

### Availability of data and materials

The raw reads for all RNA-seq data, the ATAC-seq data from the ruminal and esophageal epithelial cells and hepatocyte have been deposited at the Sequence Read Archive (SRA) under project number PRJNA485657. The customized scripts have deposited in GitHub (https://github.com/xiangyupan/CNEReg).

### Ethics approval and consent to participate

All implemented experiments were approved by the Institutional Animal Care and Use Committee and were in strict accordance with good animal practices as defined by the Northwest A&F University (protocol number: NWAFAC1008). All efforts were made to minimize animal suffering.

