## Supplementary figures for "Interpreting ruminant specific conserved non-coding elements by developmental gene regulatory network"

**^Contents^**

[Supplemental Table S4 - 1,061 active-RSCNEs are enriched for binding motifs of transcriptional regulators known to play vital role in rumen development (128 motifs with Benjamini *q*-value < 1.00×10^-3^).	11](#_Toc85704437)

### Supplemental Figures

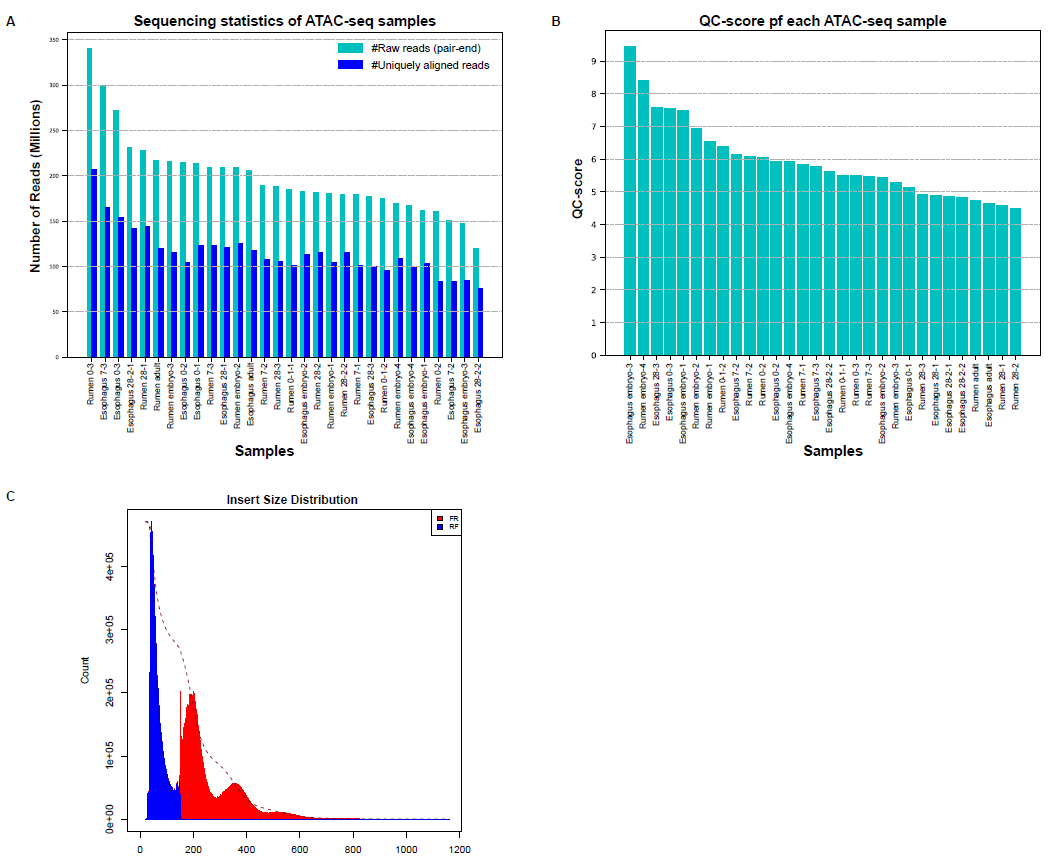

Fig. S1. Data quality check for the ATAC-seq samples by their sequence depth, fragment distribution, and QC score. (**A**) The number of raw reads and percentile of uniquely aligned reads. The number of raw reads ranges from 110 to 330 million for each ATAC-seq sample, and the percentiles of uniquely aligned reads across all samples is 59% on average. (**B**) QC scores of all ATAC-seq samples. QC score is defined as the ratio of total reads count at TSS centered up- and down-stream 2-kb window to the randomly selected background [-3k, -2k] among all genes. All the samples are required to reach the standard score 4, i.e., the fold change is large than 1. This indicates the ATAC-seq signals are enriched in open chromatin regions such as promoters. (**C**) Insert size distribution of one example esophagus-0-1. The other 29 samples show similar pattern. This fragment length distribution reveals a sharp peak at less than 100bp regions for nucleosome-free fragments and the second large peak is within 200bp for the mono nucleosome fragment. Again, this indicates good data quality.

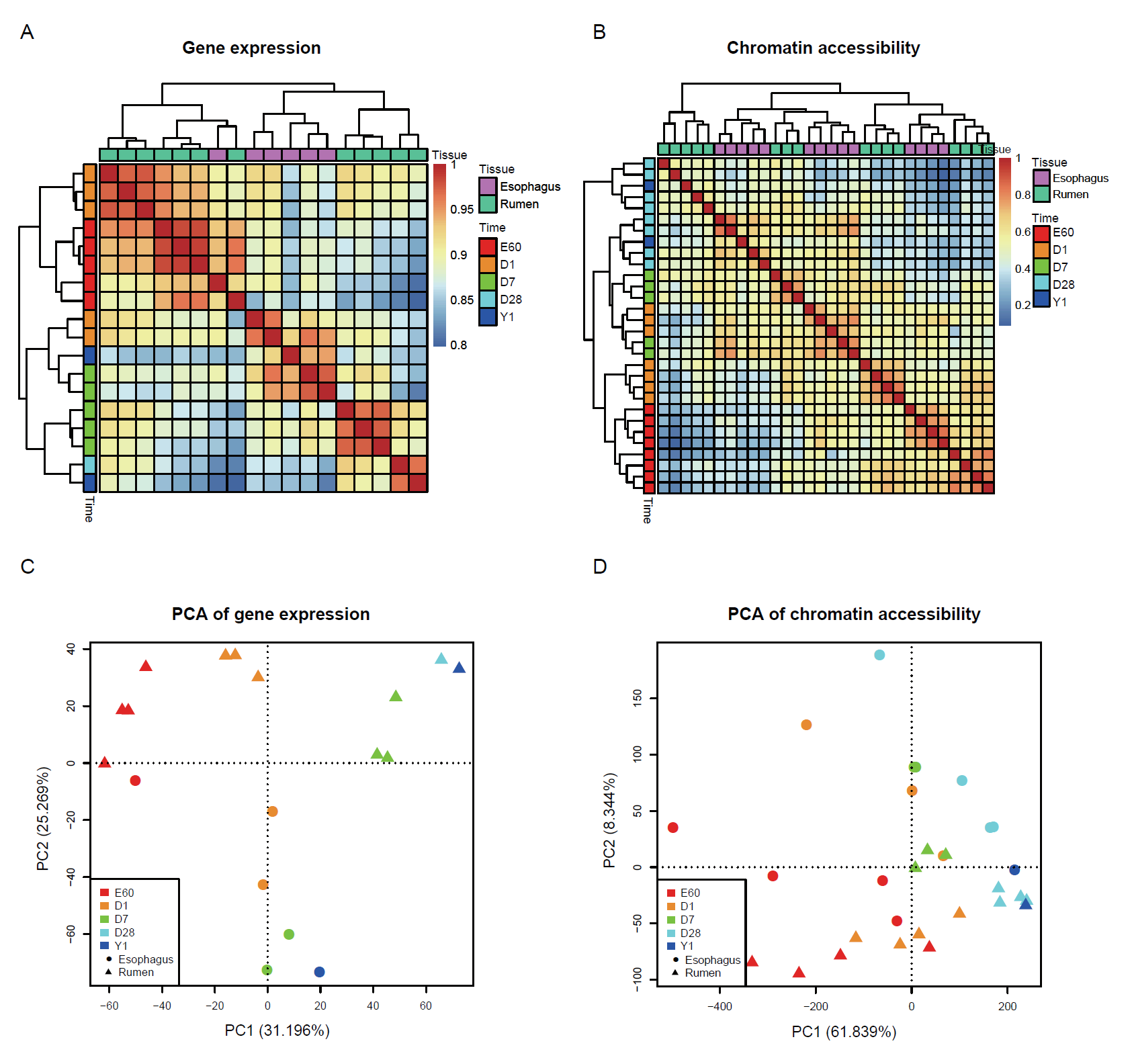

Fig. S2. Paired expression and chromatin accessibility time series data reveals the regulatory landscape for rumen and esophagus development. **(A, B)** Hierarchical clustering of gene expression and chromatin accessibility of all samples including biological and technical replicates. **(C, D)** Principal component analysis for 14,637 genes and 178,651 open chromatin regions. Rumen and esophagus show consistent developmental pattern at both gene expression and chromatin accessibility levels. The variation between rumen and esophagus is less than variation among developmental stages.

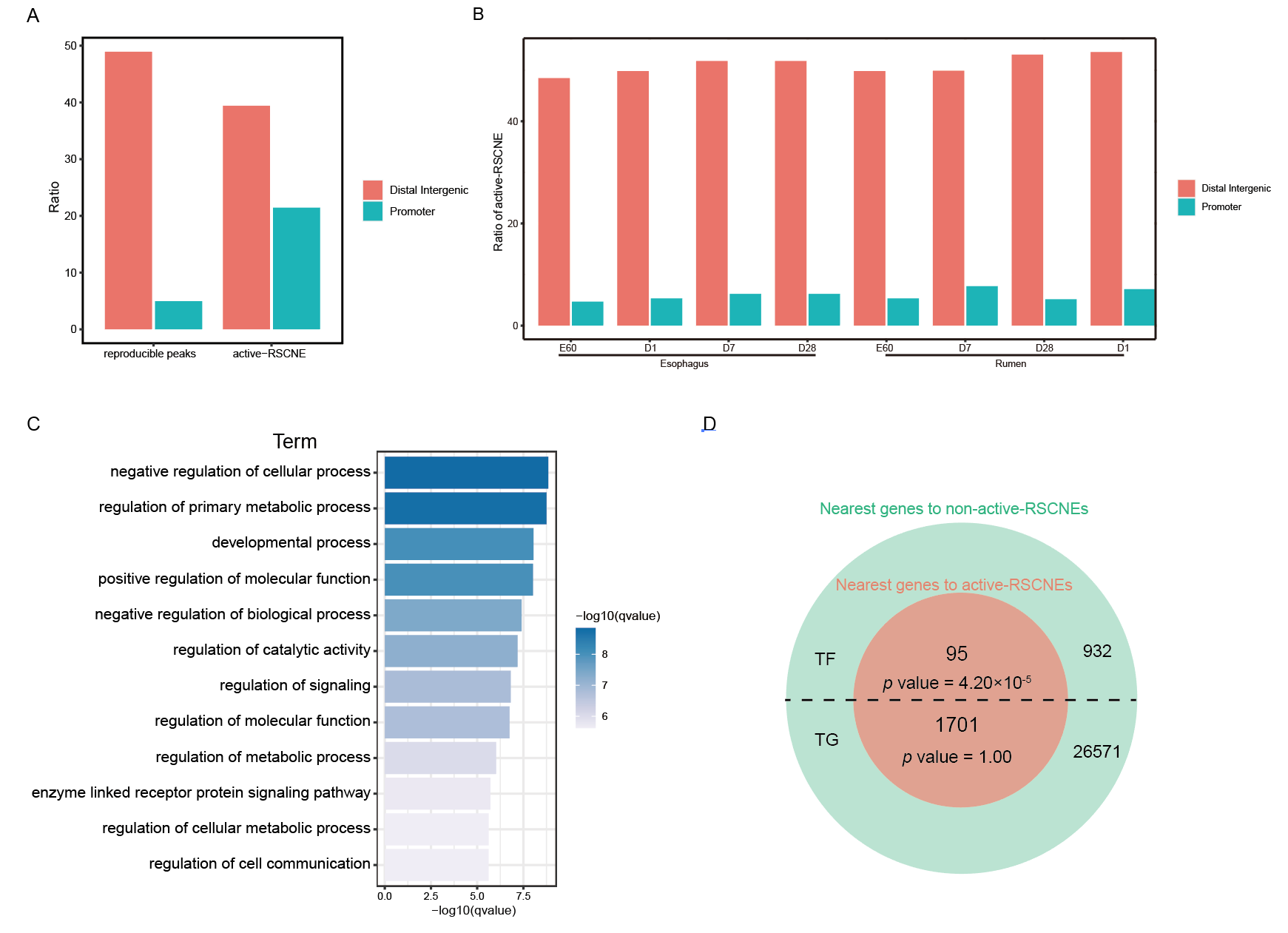

#### **Fig. S3. Further characterization of active-RSCNEs.** (A) The percentage of location in distal intergenic and promoter for reproducible peaks and active-RSCNEs. (B) The percentage of location in distal intergenic and promoter for active-RSCNEs at each developmental stage of rumen and esophagus. (C) GO enrichment analysis of active-RSCNEs’ 1,796 genes nearby. (D) Fisher’s exact test shows active-RSCNEs’ 1,796 genes nearby are enriched in TFs (*p* value = 4.20×10^-5^) but not non-TF genes (*p* value = 1.00, no significance).

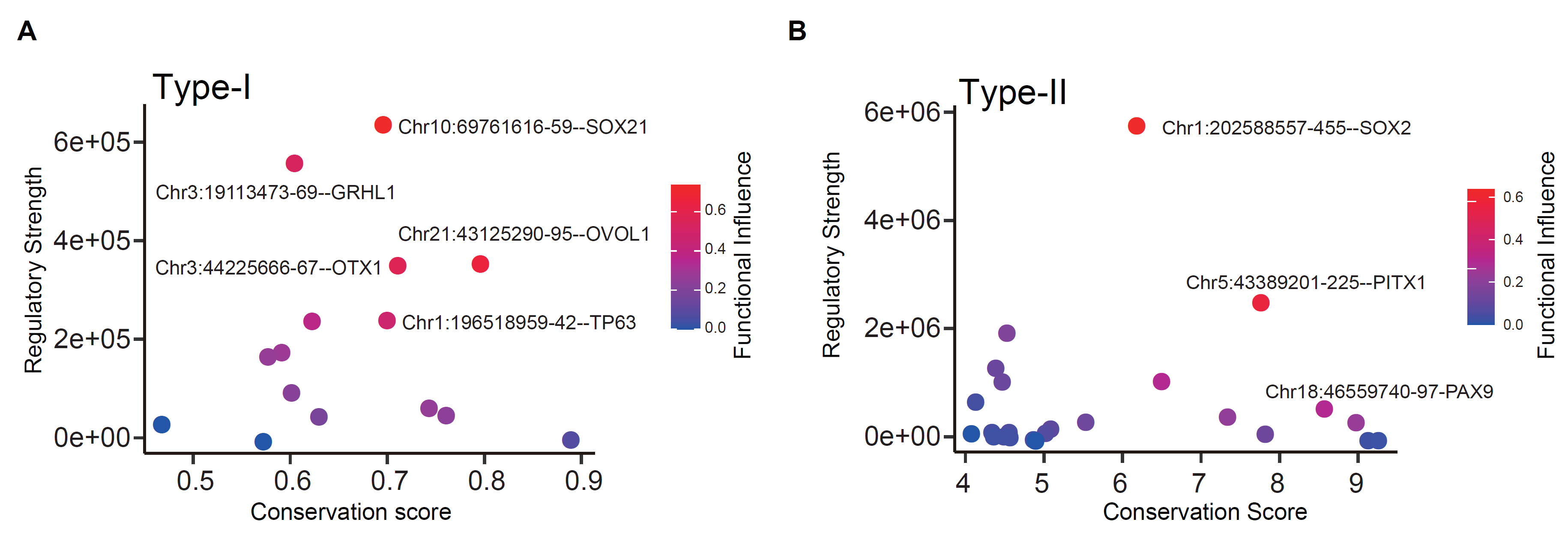

Fig. S4. Relationships between the regulatory strength and the conservation score of TTF upstream network. Scatter diagrams show the regulatory strength (y-axis), the conservation score (x-axis), and the functional influence score (node color) of Type-I (**A**) and Type-II (**B**) active-RSCNEs with regulated gene which are in the network from **Fig. 4**.

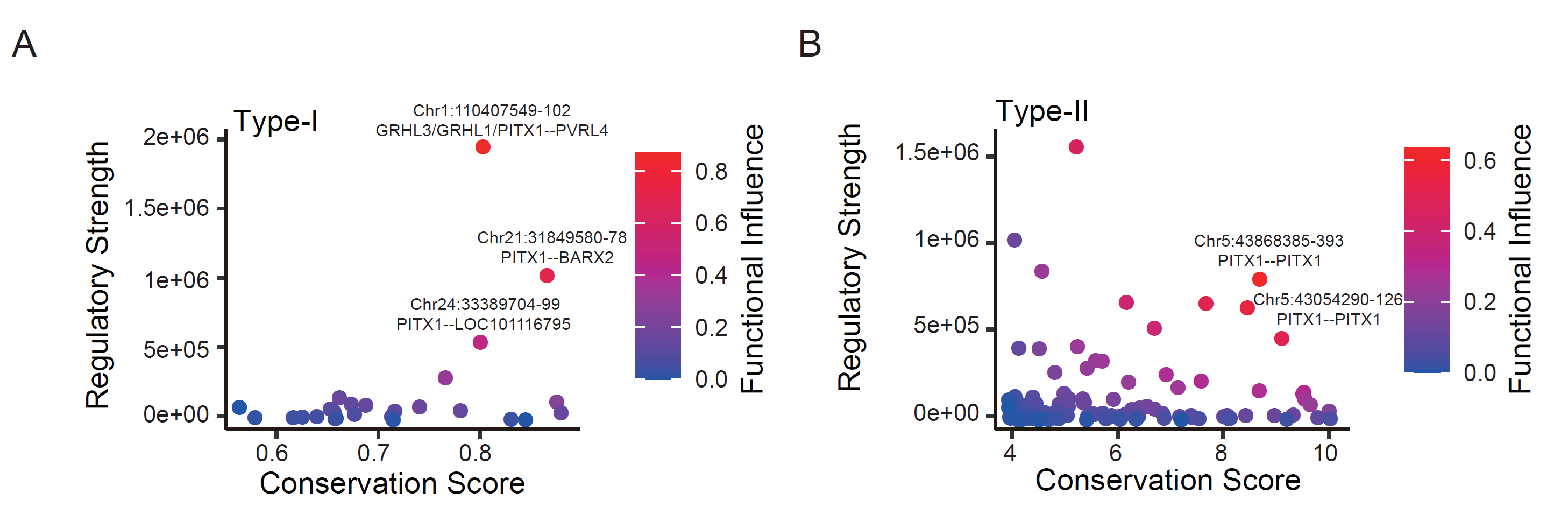

Fig. S5. Relationships between the regulatory strength and the conservation score of TTF downstream network. Scatter diagrams show the regulatory strength (y-axis), the conservation score (x-axis), and the functional influence score (node color) of Type-I (**A**) and Type-II (**B**) active-RSCNEs with regulated gene, which from the network in **Fig. 5**.

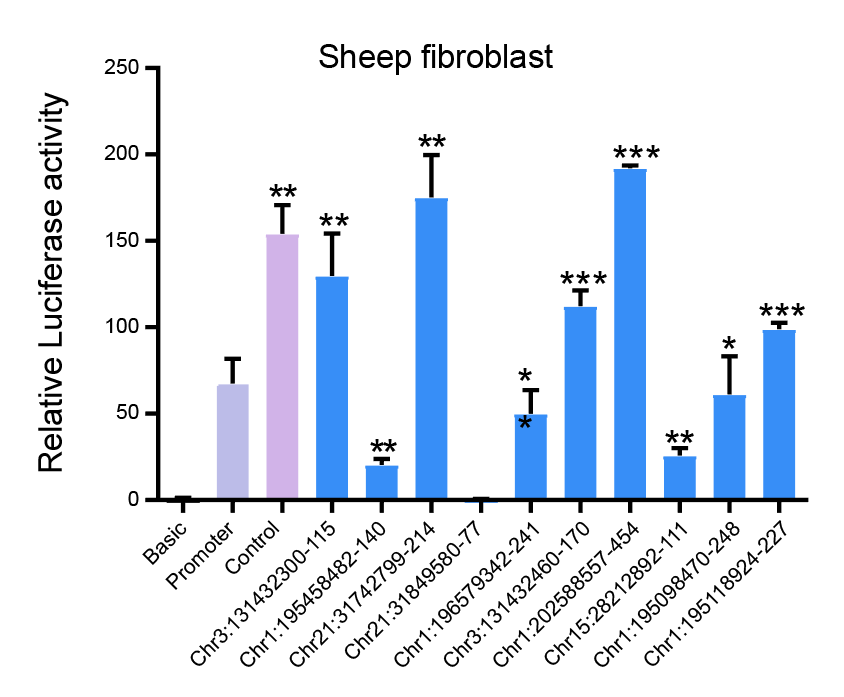

Fig. S6. Luciferase activity assays of 10 active-RSCNEs with top functional influence score. 9 of 10 show significant regulatory activity in PGL-3 promoter (one side t-test, * *p*-value<0.05, ** *p*-value <0.01, *** *p*-value<0.001).

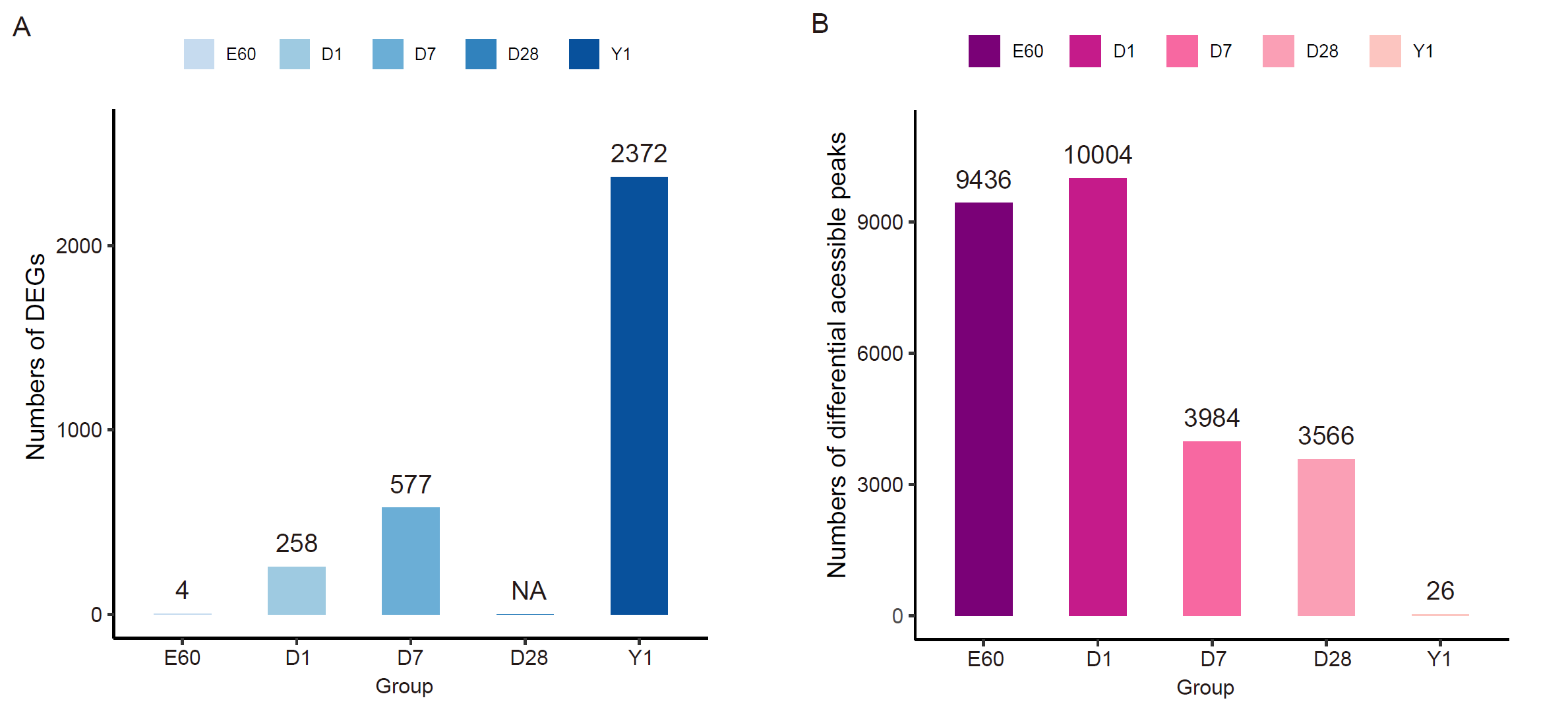

Fig. S7. Differentially expressed genes and differentially accessible peaks between rumen and esophagus at each stage. **(A)** Barplots showing the number of differentially expressed genes between rumen and esophagus at each stage. (**B**) Barplots showing the number of differentially accessible peaks between rumen and esophagus at each stage.

### Supplemental Table Descriptions

#### Supplemental Table S1 - Statistic of 37 ATAC-seq data used in this study.

#### Supplemental Table S2 - Statistic of RNA-seq data used in our study.

#### Supplemental Table S3 - 1,601 active-RSCNEs.

#### Supplemental Table S4 - 1,061 active-RSCNEs are enriched for binding motifs of transcriptional regulators known to play vital role in rumen development (128 motifs with Benjamini *q*-value < 1.00×10^-3^).

#### Supplemental Table S5 - 18 rumen toolkit TFs (TTFs).

#### Supplemental Table S6 - Upstream regulatory network of 18 rumen toolkit TFs.

#### Supplemental Table S7 - Downstream regulatory network of 18 rumen toolkit TFs.

#### Supplemental Table S8 - Differential regulatory sub-network between rumen and esophagus.

#### Supplemental Table S9 - GO enrichment analysis of 52 TGs in the differential regulatory sub-network between rumen and esophagus. BP denotes Biological Process, MF denotes Molecular Function, and CC denotes Cellular Component.

#### Supplemental Table S10 - The gene expression profile of 655 rumen specifically expressed genes (RSEGs) which showed by FKPM value.
